# Predicting Early Transitions in Respiratory Virus Infections via Critical Transient Gene Interactions

**DOI:** 10.1101/2025.04.18.649619

**Authors:** Chengshang Lyu, Anna Jiang, Ka Ho Ng, Xiaoyu Liu, Lingxi Chen

**Affiliations:** Department of Biomedical Sciences, City University of Hong Kong, Hong Kong; Department of Computer Science, City University of Hong Kong, Hong Kong; Tung Biomedical Sciences Centre, City University of Hong Kong, Hong Kong

## Abstract

Early detection of respiratory virus infections, such as influenza A (H3N2), is critical for timely intervention and disease management. Conventional biomarkers often overlook the complex and dynamic nature of gene regulatory changes, while existing predictive models frequently lack automation and robust external validation. Thus, we present CRISGI (Critical tran-Sient Gene Interaction), a computational framework that detects early-warning signals of infection by identifying dynamic changes in gene-gene interactions—termed critical transient interactions—from bulk RNA-seq data. CRISGI leverages critical transition (CT) theory to capture a GRN’s unstable intermediate state, known as the CT stage, before irreversible phenotypic shifts. Applied to a human challenge study with H3N2, CRISGI identified 128 critical transition edges (128-TER). These were used to train predictive models capable of forecasting symptom status and onset timing. 128-TER was then validated across six temporal transcriptomic datasets involving three respiratory viruses (H3N2, H1N1, HRV). The 128-TER consistently distinguished symptomatic individuals, predicted infection onset, and revealed phenotype-specific enrichment patterns. Notably, CRISGI captured immune-related transitions involving interferon-stimulated genes (e.g., IFIT1, CXCL10), underscoring their role in early host defense. CRISGI advances early-warning biomarker discovery by integrating interaction-level dynamics and predictive modeling. Its reproducibility across viruses highlights shared immune activation pathways, supporting its utility in both research and clinical contexts.

## Introduction

Detecting early warning signals for respiratory virus infections such as influenza A virus is crucial for timely clinical intervention and disease management (1). Early detection can enable preemptive therapeutic strategies, reduce transmission, and prevent progression to severe outcomes. However, current diagnostic methods often detect infection only after symptom onset or viral proliferation, leaving a critical window of intervention unaddressed (2).

Biological fate decisions during disease progression are governed by critical transitions (CTs) within complex systems such as gene regulatory networks (GRNs) (3). According to CT theory, a GRN begins in a stable *normal stage*. Upon perturbation, structural fluctuations driven by *critical transient genes* cause a spike in *critical transient signals*, marking an unstable *CT stage*. The system eventually restabilizes into a new, irreversible *phenotype stage* (3, 4). In the context of viral infection, a respiratory virus may induce a CT stage in a healthy individual, which can lead to either the development of severe symptoms or resolution into an asymptomatic state (2, 4).

Despite the promise of CT theory in revealing early-warning signals of virus infection progression (2, 4–7), several challenges remains. (i) Most methods infer criticality by quantifying increased variance and covariance among genes, yet they largely prioritize individual gene or module-level signals. This overlooks dynamic gene interactions (e.g., co-operative regulation, feedback loops) that underpin bifurcations in GRNs. (ii) The predominance of unsupervised approaches, which rank signals based solely on magnitude (e.g., top 5%) (5, 6, 8–11), often fails to isolate phenotype-specific drivers from mixed signals. (iii) Most studies fail to incorporate CT signals into predictive models capable of forecasting impending transitions or estimating onset timing; instead, they rely on manual inspection of signal patterns (e.g., peaks or trends), limiting scalability and reproducibility. (iv) CT biomarkers are typically derived from single cohorts with minimal internal or external validation, hindering generalizability and adoption in clinical practice.

To fill the gaps, we propose a computational algorithm called CRISGI (CRItical tranSient Gene Interaction). Using bulk RNA-seq data from H3N2-infected patients, CRISGI identified 128 critical transition interactions (128-TER) and built automated models to predict symptom status and onset timing. The model and biomarkers were validated across six temporal bulk RNA-seq datasets challenged with three viruses (H3N2, H1N1, HRV). Evaluation was performed across three scenarios using CRISGI scores of the 128-TER: (1) predicting post-challenge symptom status; (2) identifying onset time in symptomatic subjects; and (3) testing if phenotype- and sample-specific CRISGI enrichment associates with symptoms. In all evaluations, CRISGI demonstrated robust performance. Finally, we compare the commonality and specificity of the 128-TER signature with previously published viral infection biomarker panels, highlighting conserved immune-related pathways (IFIT1 and CXCL10) that may serve as broadly applicable early warning markers.

## Methods

### Problem formulation

Consider an observation-gene expression matrix ***X*** ∈ ℝ^*N ×M*^, where *N* denotes the number of observations and *M* represents the number of genes. Each observation corresponds to an individual sample in bulk RNA-seq data.

To model gene interaction, we define an interaction network, *G* = (𝒱, ℰ), where vertices *µ, ν* ∈ 𝒱 represent genes, and edges (*µ, ν*) ∈ ℰ represent their interactions. Importantly, the structure of this network varies across observations. In time-series applications, the network may exhibit stability initially, undergo dynamic fluctuations due to external perturbations, and eventually return to equilibrium. In CRISGI, we take the earliest observations as a reference representing the initial stable condition. For subsequent observations, we propose an entropy-based transition score to quantify the perturbation level relative to the reference. This statistical perturbation analysis allows us to characterize whether the system is stable or perturbed over time.

Observations may also represent distinct critical states or phenotypes, such as symptom status after a viral infection. In CRISGI, users can designate one phenotype as the perturbation group and others as reference groups. Transition scores are then subjected to statistical tests to identify CT interaction dynamics. This analysis reveals differentially expressed interactions (DERs) and trend expressed interactions (TERs) within the gene network that are specific to phenotypes and time progression, respectively. After deriving TERs, CRISGI employs a convolutional neural network (CNN) to predict the occurrence of CT stage based on incoming time-ordered observations. If a CT stage is detected, CRISGI applies a shortterm energy detection method to estimate the onset time of the CT stage.

### Background gene interaction network topology

We utilized a well-validated protein-protein interaction (PPI) network as the background for the gene interaction modeling. PPIs were sourced from the STRING database (version 12, https://www.string-db.org, `9606.protein.links.v12.0.txt.gz`) (12), applying an interaction score threshold. Protein stable IDs from STRING were mapped to gene names using BioMart (https://www.ensembl.org/biomart/martview/, Ensembl Genes 112, human genes GRCh38.p14). For each observation-gene expression matrix ***X***, the background network *G* was further filtered using high-variable genes (HVGs).

### Interaction entropy within an observation population

For the graph *G* with gene expression ***X*** carrying *N* observations, we aim to compute the entropy 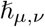 for each interaction (*µ, ν*) ∈ ℰ in following steps.

First, we use the covariance matrix as the affinity score *a*_*µ*,*ν*_ for the interaction between genes *µ* and *ν*:

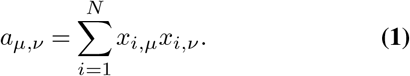

Second, we determine the interaction probability *p*_*µ*,*ν*_ between genes *µ* and *ν*:

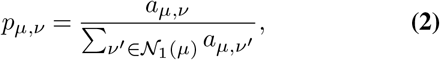

where *ν*^*′*^ ∈ 𝒩_1_(*µ*) represents the 1st-order neighbors of gene *µ*.

Third, we compute the entropy 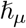 of gene *µ* by:

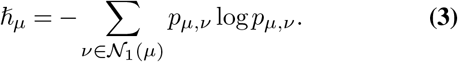

Meanwhile, with average expression 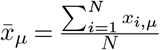 of gene *µ*, we calculate the standard deviation of gene *µ*,

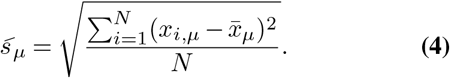

Finally, we calculate the entropy 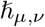 and standard deviation 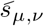 for the interaction (*µ, ν*) as the average entropy and standard deviation of gene *µ* and *ν*,

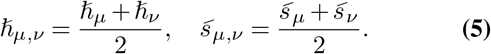

### Interaction entropy for reference and test observation

For time-series data, the first *N* ^*′*^ observations are designated as the reference population ℛ = {*i* |1 ≤ *i ≤ N* ^*′*^}, representing the initial state of the entire population. The entropy and standard deviation of each interaction (*µ, ν*) ∈ ℰ within the reference population are denoted as 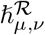 and 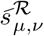, respectively.

After defining the reference population, the remaining observations—or all observations, depending on the analysis—are treated as test observations in CRISGI. For each test observation *j*, we extend the reference population to form a “test population” 𝒯_*j*_ = {*i, j* | 1 ≤ *i* ≤ *N* ^*′*^}. The entropy and standard deviation of the interaction (*µ, ν*) ∈ ℰ within the test population 𝒯_*j*_ are calculated as 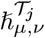 and 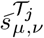, respectively.

### Transition score of test observation

We define the *transition score* for each test observation based on statistical perturbation analysis relative to the reference population. Specifically, the transition score 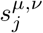 for interaction (*µ, ν*) ∈ ℰ at test observation *j* is calculated as the product of the absolute differences in entropy and standard deviation between the test population 𝒯_*j*_ and the reference set ℛ:

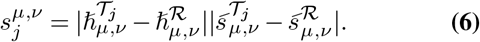

This score quantifies the perturbation of the gene interaction (*µ, ν*) at test observation *j* relative to the reference ℛ. A transition score 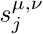 close to zero indicates that the interaction (*µ, ν*) remains stable in the test observation *j* compared to the reference. Conversely, a significantly higher 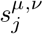 reflects substantial changes in the interaction (*µ, ν*) at observation *j*.

### Transition curve of test observation along time

For a gene interaction (*µ, ν*) within a graph, the *transition curve* ***s***^*µ*,*ν*^ represents the transition scores across all *N* observations when each observation is treated as the test case against the reference ℛ, ordered chronologically:

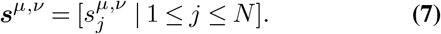

The transition curve ***s***^*µ*,*ν*^ begins at zero. If it remains near zero over time, this suggests an absence of transition dynamics in the interaction (*µ, ν*). Conversely, a pronounced upward trend in ***s***^*µ*,*ν*^ indicates significant transition dynamics, implying that the interaction (*µ, ν*) plays a critical role in time progression.

### Phenotype-specific CT interactions: DERs and TERs

Suppose these observations exhibit *K* critical statuses (phenotypes). Select one critical status as the perturbation status and the remaining statuses as reference statuses. Then, the observations belonging to the perturbation status form a perturbation group 𝒫, and the observations from the *K* − 1 reference statuses form reference groups 𝒮 = {𝒮_1_, 𝒮_2_, …, 𝒮_*K*−1_}.

In CRISGI, we aim to identify the perturbation-specific differentially expressed interactions (DERs) against all reference groups. For each interaction (*µ, ν*), we calculate the log2 fold change of transition scores between each perturbation-reference group pair (𝒫, 𝒮_*k*_) and compute test statistics, such as z-scores, p-values, and adjusted p-values. Both the Wilcoxon test (13) and t-test (14) are offered. All significant (typically with a p-value or adjusted p-value less than 0.05) and upregulated (with a log2 fold change larger than a positive threshold) interactions (*µ, ν*) form the perturbation-specific DER network against reference group 𝒮_*k*_, denoted as 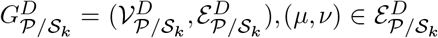. By iterating over all perturbation-reference group pairs, for interaction (*µ, ν*), we obtain the perturbation-specific DER networks against the *K* − 1 reference groups. We use the interactions (*µ, ν*) that belong to all DER networks to form the final perturbation-specific DER network against all reference groups 𝒮:

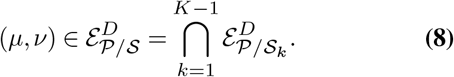

If the observations have a temporal order, given a perturbation-specific DER network, we can further identify the trend expressed interactions (TERs) network 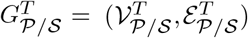 based on two conditions: (1) The transition curve ***s*** of interaction (*µ, ν*) in the perturbed group shows a significant increasing trend, determined using the Mann-Kendall test (15, 16). (2) The transition curve ***s***^*µ*,*ν*^ of interaction (*µ, ν*) in the reference groups remains constant at zero, determined using a one-sample t-test (14).

### Predicting the existence of CT stage for time-ordered observations

Given a list of observations ordered by time, for example, the samplings from the patient subject given by different timepoints. We have screened out the TER network for a specific critical status. Here, we aim to predict the existence of CT stage using the TER network for incoming observations.

First, we transfer the transition score of TER network into a heatmap image, with columns and rows stand for the TERs and observations ordered in time, respectively. Next, we applied CNN-autoencoder to the images, and connected to a multiple-layer perceptron (MLP) for a binary classifier of the existence of critical status. The structure of the CNN-autoencoder is as follows:

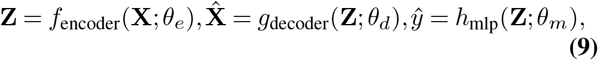

where **X** is the input image, **Z** represents the features extracted from the **X** by the encoder, and *θ*_*e*_ represents the parameters of the encoder. 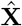 is the reconstructed image, and *θ*_*d*_ represents the parameters of the decoder. In the structure of the MLP for binary classification, *ŷ* is the predicted probability of the positive class, and *θ*_*m*_ represents the parameters of the MLP.

The loss function includes the image reconstruction loss plus the binary cross-entropy loss,

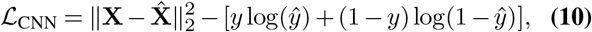

where **X** is the original image and 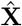 is the reconstructed image. *y* is the true label (0 or 1), and *ŷ* is the predicted probability of the positive class.

### The onset point of CT stage for time-ordered observations

To identify the onset point when a subject enters a critical status based on the TER transition curves over time, we treat this as a speech start-point detection problem. Here, each TER transition curve 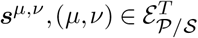 is considered a speech signal. Our goal is to detect the concurrent starting point across multiple TER signals, which have similar increasing trends but contain some noise. We employ an energy-based method, observing that TER signals within a critical status typically exhibit higher energy compared to background noise.

The first step is short-term energy extraction. We divide each TER signal into overlapping frames of a specified window size with a given overlap size. The short-term energy for a window starting at observation *i* with window length *L* is calculated by summing the squared signal values within the window:

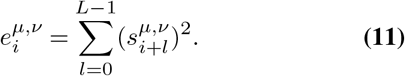

We normalize the energy values for each TER signal to have a mean of 0 and a standard deviation of 1. Summing the normalized energies across all TER signals enhances common TER activity. To further reduce noise, we apply median filtering with a kernel size of 5 to the summed energy.

We assume the smoothed TER energy follows a normal distribution, we set a threshold at the mean plus 2 standard deviations (*mean* +2*σ*) of the smoothed energy. This threshold is based on the fact that in a normal distribution, approximately 95% of the data points fall within 2 standard deviations of the mean. Values exceeding this threshold indicate significant changes in energy levels, likely corresponding to the onset of critical status. Thus, the first timepoint where the smoothed energy exceeds the *mean* + 2*σ* threshold is considered the critical status onset point.

## Results

### Detecting viral CT interactions from bulk RNA-seq dataset GSE30550

Understanding host transcriptional responses to respiratory viruses is key to elucidating virus-induced immunopathology. Using the temporal bulk RNA-seq dataset GSE30550 (influenza A H3N2) (1), we demonstrate that CRISGI can identify CT interactions during virus infection

We downloaded influenza A H3N2 dataset GSE30550 (1) from GEO. 17 healthy humans were injected with the living H3N2 virus, and their gene expression data were collected from blood at sixteen time points over 132 hours (-24, 0, 5, 12, 29, 36, 45, 53, 60, 69, 77, 84, 93, 101, 108 hours). After injecting the virus at 0 hours, 9 of 17 subjects developed flu symptoms (subjects 1, 5, 6, 7, 8, 10, 12, 13, 15), and the rest remained asymptomatic (subjects 2, 3, 4, 9, 11, 14, 16, 17). Blood taken 24 hours prior to inoculation with virus as regarded as baseline, immediately prior to inoculation as pre-challenge, and at set intervals 8 hour intervals following challenge are regard as post-challenge.

The top 5,000 highly variable genes (HVGs) were used to construct a background network with 15,753 gene interactions based on STRING database (12) with STRINGDB score > 850. The CRISGI score for each gene interaction was calculated for all samples, with all baseline samples serving as the reference group, representing a healthy state unaffected by the virus. We identified 3,441 DERs with the following criteria: the CRISGI score in the symptomatic group (n=9) is higher than in the asymptomatic group (n=8) (log2 fold change > 1 and adjusted p-value < 0.05, Wilcoxon test). We screened out 2,519 TERs conditioned on the symptomatic group shows a significant trend (p < 0.05, Mann-Kendall test), and the asymptomatic group remains constant at zero (p < 0.05, one-sample t-test) (Figure 2A-C). In Figure 2D, TER CRISGI scores significantly differed between symptomatic and asymptomatic groups. A sharp increase in CRISGI score marked a CT preceding symptom onset. All nine symptomatic subjects showed CT signals, with seven exhibiting before symptoms. No CT signals were observed in asymptomatic subjects.

**Fig. 1.**
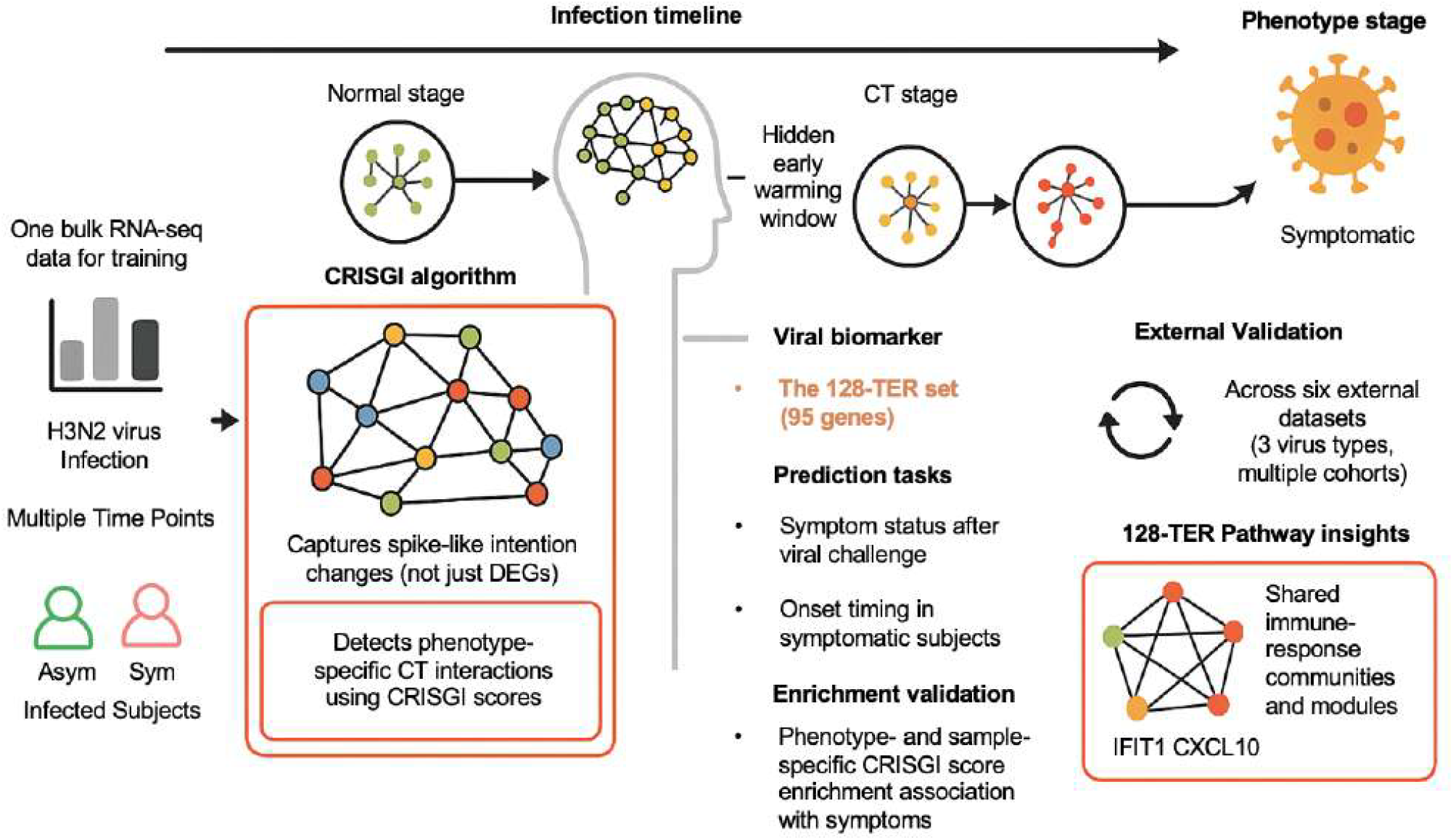
Graphic summary of the study design.

**Fig. 2.**
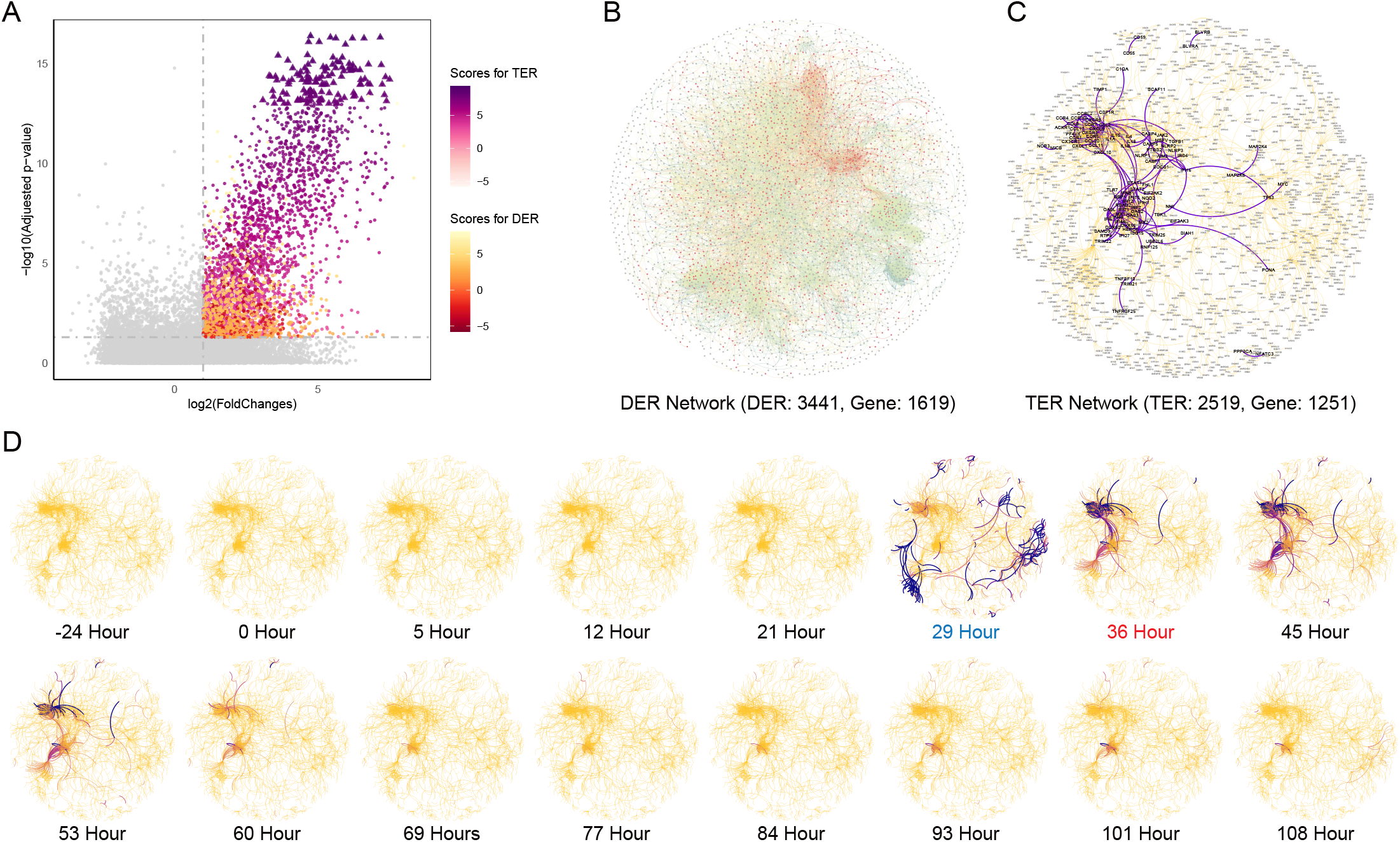
Detecting CT interactions in viral respiratory infections dataset GSE30550. **A** Volcano plot showing differentially expressed relations (DERs, color from red to yellow) and trend expressed interactions (TERs, color from pink to purple) used in the screening process. The top 5% (128) TERs are marked as triangles in the volcano plot. **B** Network plot illustrating all 2519 TERs (1251 red nodes) filtered from a total of 3441 DERs. **C** Network plot showing the 128 top TERs (marked with purple color) filtered out from all 2,519 TERs. **D** Dynamics of TERs’ CRISGI scores over challenge time for the symptomatic subject 1.

### Screening 128-TER for viral respiratory infection from GSE30550

CRISGI-based pathway enrichment on incrementally ranked top-n TERs (n = 10 to 2500, step = 10) revealed consistent enrichment in pathways relevant to Influenza A infection, including defense response to virus, NOD-like receptor signaling, cytokine and chemokine signaling, inflammatory response, and interferon responses. The overall decline in nTopRatio with increasing *n* suggests that the most biologically relevant pathways are concentrated among the top-ranked TERs. Therefore, we focused on the top 5% (n = 128) TERs (Figure 3A), a threshold widely used in prior studies (5, 6, 8–11). Comparing the 128-TER (purple in Figure 2C) with sample-specific top 5% scoring TERs (dark purple in Figure 2D), we found near zero overlap in pre-onset stages, which sharply increased before onset, averaging 21.34% post-onset (orange border in Figure 3B). In contrast, overlaps in asymptomatic subjects remained near zero (avg. 0.54%). Furthermore, enrichment analysis of the 128-TER echoed results from the full-ranked nTopRatio analysis (Figure 3C).

**Fig. 3.**
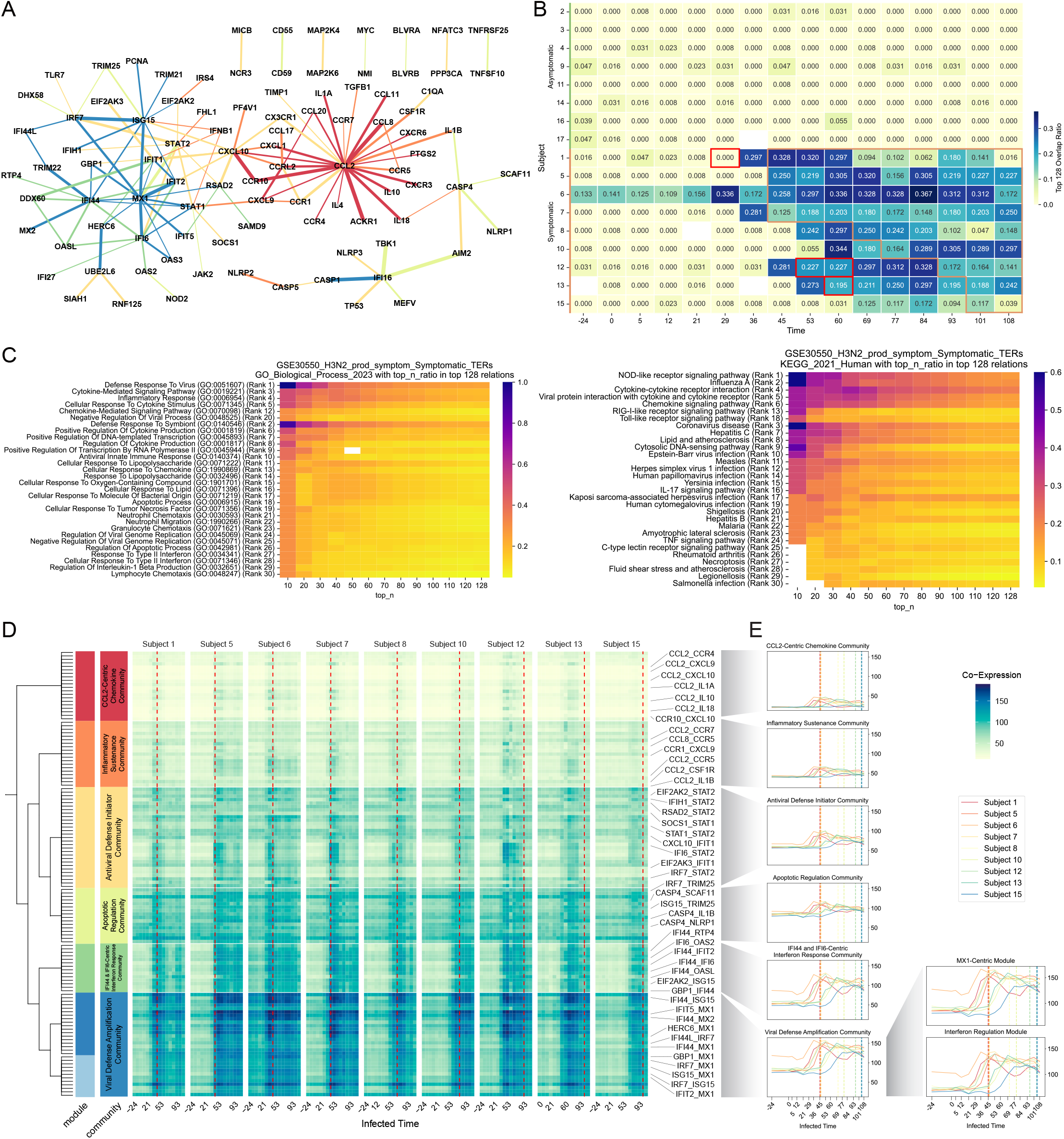
128-TER as CT interactions in viral respiratory infections dataset GSE30550. **A** The top 128-TER networks with specific colors corresponding to the SEAT clustering results in Figure 3D. **B** Heatmap of overlap ratio among all subjects. The yellow zone stands for the post-onset stages, and the red zones stand for special alterations in the pre-onset stages. **C** Enrichment analysis for the top 5% TERs in KEGG and GOBP Pathways, presented in increasing order of the accumulated “nTopRatio”. **D** The clustering results and co-expression heatmap of 128-TER processed by SEAT algorithm, containing 6 communities and 7 modules. **E** Line charts of the average co-expression scores of all communities/modules. KEGG: Kyoto Encyclopedia of Genes and Genomes. GOBP: Gene Ontology Biological Process.

Next, we examined the co-expression dynamics of the 128-TER across nine symptomatic subjects. In Figure 3D–E, these TERs exhibit a consistent pattern: stable expression followed by a sharp increase at or before symptom onset (red dashed line), supporting the role of CRISGI scores in capturing CTs.

Despite this shared trend, notable heterogeneity exists. To explore this, we built an interaction hierarchy comprising six communities with distinct co-expression patterns and potential regulatory functions (Figure 3D-E). The CCL2-centric chemokine community 0 is central to recruiting immune cells, balancing inflammation, and coordinating antiviral defense. The inflammatory sustenance community 1 focuses on sustaining chronic inflammation and enhancing antiviral responses. The antiviral defense initiator community 2, centered around STAT1 and STAT2, triggers early interferon responses to fight viral infections. The apoptotic regulation community 3 regulates immune homeostasis via apoptosis and pyroptosis to eliminate infected cells. The IFI44 and IFI6-centric interferon response community 4 amplifies interferon signaling for efficient viral clearance. Finally, the viral defense amplification community 5 plays a key role in long-term immune activation, particularly through antiviral genes like MX1 and ISG15, which sustain the immune response after symptom onset. These six communities work together to ensure a coordinated and efficient immune response to infections while preventing excessive tissue damage.

Lastly, while most symptomatic subjects exhibit viral response activity within the 128-TER, distinct sample-specific TER signals emerge earlier in some cases. Notably, subject 1 at 29h, subject 12 at 53h and 60h, and subject 13 at 60h display significant TERs outside the 128 set (blue markers in Figure 2D). As highlighted in red boxes in Figure 3B, subject 1–29 shows zero overlap with the 128-TER. Subject 12’s overlap fluctuates: starting at 28.1% (45h), decreasing slightly at 53h and 60h (22.7%), and rising again to 29.7% (69h). Subject 13’s overlap drops from 27.3% to 19.5% at 60h. These unique signals involve apolipoproteins and kinases (subject 1–29), Fanconi anemia and influenza-related genes (subject 12–53), pathways linked to cell cycle, calcium signaling, and immune response (subject 12–60), and lipid metabolism, interferon, and inflammation (subject 13–60).

### 128-TER act as viral CT biomarkers for GSE30550 dataset

We assessed the 128-TER as viral CT biomarkers for GSE30550. Evaluation was performed across three scenarios using CRISGI scores of the 128-TER: (1) predicting post-challenge symptom status; (2) identifying onset time in symptomatic subjects; and (3) testing if phenotype- and sample-specific CRISGI enrichment associates with symptoms.

In the first scenario, we generated biomarker heatmaps for each subject (Figure 4A). Using these heatmaps and symptom labels from 17 subjects, we trained a three-layer CNN (3L-CNN) autoencoder to classify subjects as symptomatic or asymptomatic. One-layer CNN (1L-CNN) and logistic regression (LR) served as baselines. All models achieved perfect performance with 100% accuracy and AUC.

**Fig. 4.**
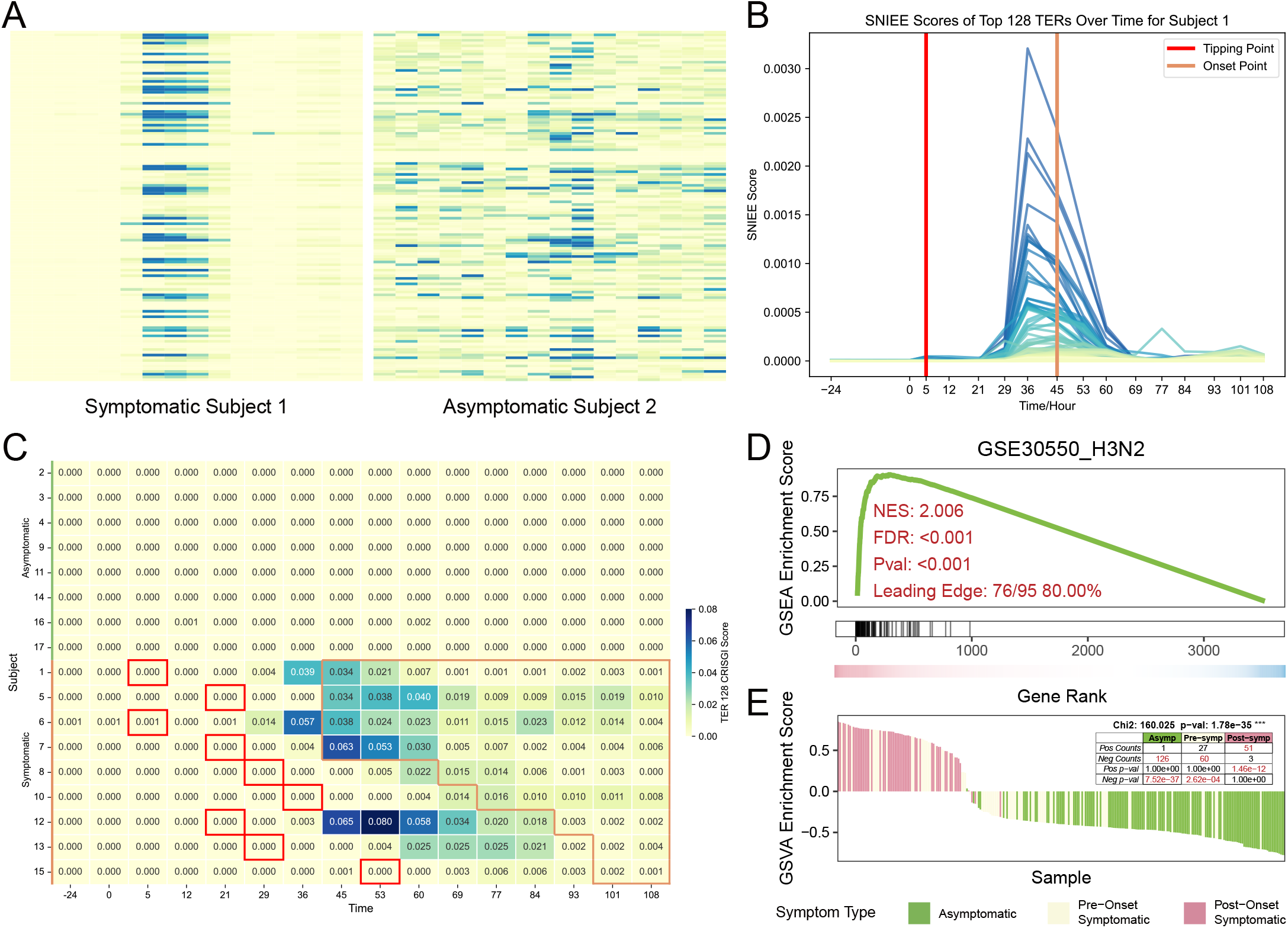
128-TER act as viral CT biomarkers for GSE30550 dataset. **A** Heatmap of 128-TER’s CRISGI scores comparing the symptomatic subject 1 and the asymptomatic subject 2 during training, with rows as TERs, columns as time points. **B** Short-energy plot showing the predicted onset time (29 hours, red line) and actual clinical symptom onset (45 hours, orange line) for the symptomatic subject 1. **C** Heatmap of CRISGI scores for the 128-TER, with rows representing subjects and columns denoting time points. Each cell displays the overlap ratio between the 128-TER and the sample-specific top 5% TERs. Red boxes indicate inferred onset times, while orange borders mark actual clinical symptom onset. **D** Phenotype-specific CRISGI enrichment plot of 128-TER as target gene set for GSE30550 dataset. **E** Sample-specific CRISGI enrichment plot of 128-TER as target gene set for GSE30550 dataset.

In the second scenario, we transformed 128 TER CRISGI scores over time into short-term energy signals (Figure 4B) and applied an energy detection algorithm to predict symptom onset in symptomatic subjects. The method identified onset accurately for all 9 subjects, with an average lead time of 44.56 hours before clinical symptoms onset (Figure 4C). CRISGI clearly infers the onset time (red boxes) for all 9 symptomatic subjects prior to their actual clinical symptom onset (orange borders).

In the third scenario, we performed both phenotype-specific and sample-specific CRISGI enrichment analyses. For phenotype-specific enrichment, we ranked all gene interactions by the average z-scores from differential expression analysis between symptomatic and asymptomatic samples. Using the genes within the 128-TER as the target set, we applied gene set enrichment analysis (GSEA) (17). A significantly positive enrichment score indicates that the 128-TER exhibit marked fluctuations in symptomatic samples compared to asymptomatic ones, reflecting a strong association with the symptomatic phenotype. For sample-specific enrichment, we ranked gene interactions by their CRISGI scores for each individual sample across time points and applied gene set variation analysis (GSVA) (18). A significantly positive enrichment score suggests that the 128-TER are actively fluctuating in that sample, reflecting dynamic transcriptomic shifts related to infection response. As a result, the phenotype-specific CRISGI enrichment yielded a strong association between the 128-TER and the symptomatic phenotype (NES = 2.0064, FDR < 0.001; Figure 4D), with 80% of hit genes in the leading-edge subset. Sample-specific CRISGI enrichment revealed significant activation of 128-TER in post-onset symptomatic samples (n = 51 with positive ES, p = 1.46e–12), while pre-onset symptomatic (n = 60 with negative ES, p = 2.62e–04) and asymptomatic samples (n = 126 with negative ES, p = 7.52e–37) showed consistent suppression (Figure 4E).

### External validation of 128-TER as viral CT biomarkers across six bulk RNA-seq datasets

Next, we validated whether CRISGI scores of 128-TER can act as CT biomarkers across six external bulk RNA-seq datasets. GSE52428_H3N2 (n=17) (19), GSE73072_H3N2_DEE2 (n=17) (20), GSE52428_H1N1 (n=24) (19), GSE73072_H1N1_DEE3 (n=24) (20), GSE73072_H1N1_DEE4 (n=19) (20), and GSE73072_HRV_DUKE (n=27) (20) were downloaded. Peripheral blood gene expression data were collected from each subject at baseline, pre-challenge, and post-challenge time points. Post-inoculation, some subjects developed symptoms while others remained asymptomatic. In Figure 5A, the complete representation of all 128 TERs across the six validation datasets supports their appropriateness for validation, as no TERs are missing.

**Fig. 5.**
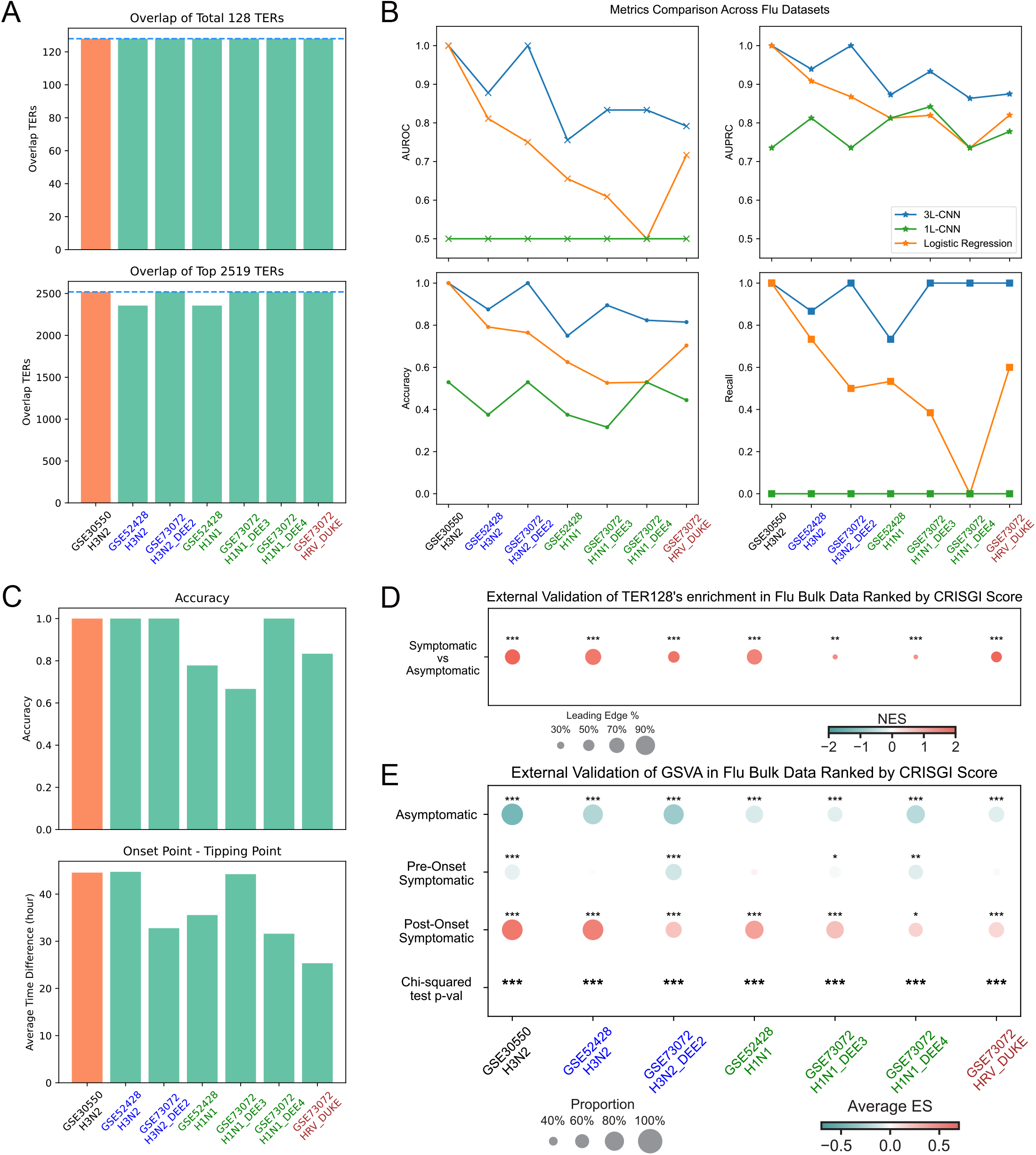
External validation of 128-TER as CT biomarkers for viral respiratory infections. **A** The number of TERs present from the 128-TER and 2,519-TER sets across the six validation datasets. **B** Training and validation performance in predicting post-challenge symptom status. **C** Training and validation performance in detecting onset time of symptomatic subjects. **D** Phenotype-specific CRISGI enrichment plot of 128-TER as target gene set across six validation datasets. **E** Sample-specific CRISGI enrichment plot of 128-TER as target gene set across six validation datasets.

For each validation dataset, we used pre-challenge and baseline samples as references to compute CRISGI scores of 128 TERs for all samples. These scores were then used to predict symptom status and detect onset times across the six validation datasets.

In Figure 5B, the 3L-CNN classifier outperformed both the 1L-CNN and logistic regression baselines. Specifically, the 3L-CNN achieved perfect performance on the GSE72072_H3N2_DEE2 dataset, with flawless AUCROC, AUPRC, accuracy, specificity, sensitivity, and precision. It also performed strongly on the other H3N2 dataset, GSE52428. For H1N1 and HRV infections, the classifier maintained robust performance, achieving approximately 80% AUROC and accuracy, around 90% AUPRC, and near-perfect recall, demonstrating strong generalizability across different viral types.

Onset time detection (Figure 5C) shows 100% early-warning success for symptomatic subjects in two external H3N2 datasets, with CT signals appearing 44.72h and 32.78h before actual symptom onset. For H1N1, early-warning was successful in 67–100% of cases, with lead times of 31.6–44.22h. For HRV, 83% of subjects were accurately early-warned, with an average lead time of 25.33h.

Finally, enrichment analysis (Figure 5D-E) further supports the positive association between 128-TER and symptoms: all six datasets show significantly positive symptomatic-specific NES values, and sample-specific ES values are significantly enriched in post-onset symptomatic samples.

### Commonality and specificity of 128-TER with known viral symptom-related panels

In this section, we examine the commonality and specificity between the 95 genes within the 128-TER and established viral symptom-related gene panels. A total of 13 panels were curated across four datasets (GSE30550_H3N2, GSE52428_H3N2, GSE52428_H1N1, and GSE17156_HRV), each derived using distinct algorithms and methodological approaches. The overlap between the 128-TER genes and known panel genes from the GSE30550_H3N2 dataset ranges from 32 to 4 genes. In comparison, overlaps with known panel genes from other H3N2 datasets and different virus datasets range from 16 to 6 genes, indicating reduced commonality as the dataset and virus type diverge (Figure 6A). From another perspective, the 128-TER genes share 55%–44% and 23%–1% overlap with known panels from GSE30550_H3N2, exhibiting a decreasing trend as the size of the reference panel increases (Figure 6B).

**Fig. 6.**
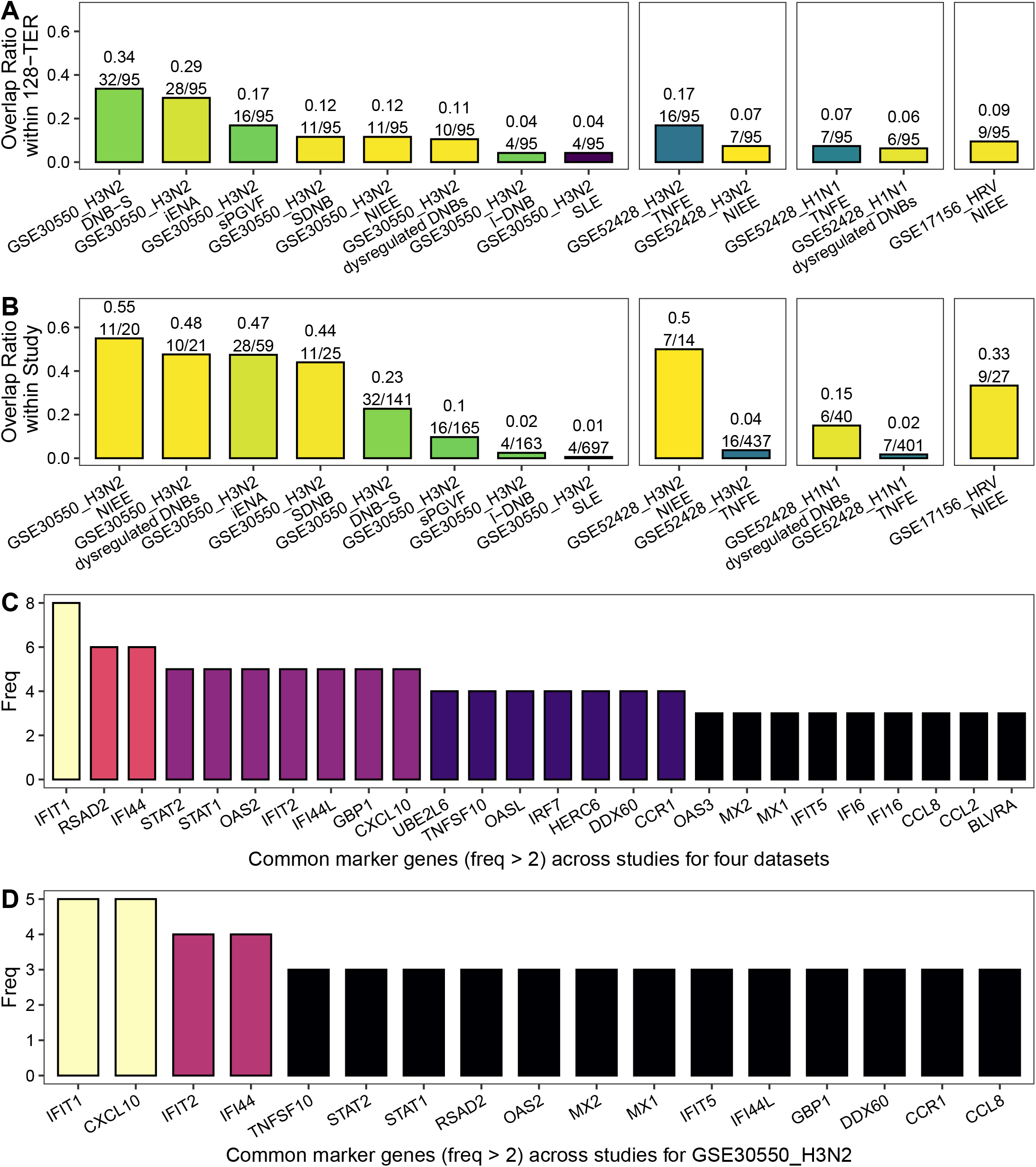
Commonality of 128-TER with other viral CT biomarker panels. **A** Overlap ratio calculated as the number of shared genes divided by the 95 genes in 128-TER, compared against those from 13 other studies across four virus infection datasets. **B** Overlap ratio calculated as the number of shared genes divided by the marker genes identified in the 13 other studies across four virus infection datasets. **C** The common 128-TER shared marker genes (freq > 2) across 13 studies for four datasets. **D** The common 128-TER shared marker genes (freq > 2) across 8 studies for GSE30550 dataset.

These overlapping genes are predominantly immune-related, including interferon-induced/related genes (IFIT1, IFIT22, IFI44, RSAD2, MX1, MX2), chemokine-related genes (CCR5, CCR1, CCL2, CCL8, CXCL10), and other antiviral genes (STAT1, STAT2, OAS2, OASL). Notably, IFIT1 is the most frequently recurring gene, appearing in 8 out of 13 known panels across all four datasets (Figure 6C). Focusing specifically on the eight GSE30550_H3N2 panels, both CXCL10 and IFIT1 are the most prominent, each appearing in five times (Figure 6D).

Among the top recurrent overlapping genes, IFIT1 and IFIT2 encode proteins acting as sensors of viral single-stranded RNAs and inhibiting viral mRNA expression (21, 22). CXCL10, a chemokine induced by interferon-gamma, is crucial for recruiting T cells and NK cells to sites of inflammation (21, 22). RSAD2 has been confirmed as an essential antiviral factor inhibiting influenza infection (23), and IFI44 plays a key role in controlling RSV infection (24). Additionally, each top recurrent overlapping gene belongs to multiple gene interaction communities (Figure 3E), suggesting a central regulatory role (as a *connector hub*) in coordinating diverse functional modules involved in antiviral responses. This connector role may explain their repeated identification across different gene panels, regardless of dataset, virus type, or analytical approach.

Among the 44 unique genes in the 128-TER, the majority are immune-related and are enriched in pathways and terms such as cytokine-cytokine receptor interaction, immune response, and viral protein interaction with cytokine and cytokine receptors. Moreover, these unique genes emphasize the regulation of apoptosis, inflammation, and the innate immune response, suggesting a comprehensive role in orchestrating the body’s defense mechanisms.

When comparing the panels across various algorithms and datasets, the overlap percentages vary significantly, ranging from 1% to 55%. This variability underscores the influence of methodological differences and dataset- and virus-specific factors on the identification of biomarkers. Despite these fluctuations, the core immune-related genes captured by the 128-TER consistently showed in known panels. In all, the 128-TER demonstrates its robustness and serves as a valuable resource for understanding the molecular mechanisms underlying immune responses in viral respiratory infections, offering both common and unique insights that complement previous studies.

## Discussion

Timely detection of viral infections remains a global priority, especially for respiratory viruses capable of rapid spread and severe disease. While many studies focus on individual gene or gene module dynamics with unsupervised approaches, our study highlights the predictive value of interaction-level dynamics within gene regulatory networks—capturing the instability and complexity that precede specific phenotypic outcomes.

CRISGI introduces a critical shift in early-warning biomarker discovery by focusing on critical transient interactions that signal a GRN’s transition from stability to bifurcation. This aligns with CT theory, where early signals arise not from static changes but from increased system fluctuations due to cooperative gene behavior. Our findings demonstrate that these dynamic features, when properly extracted and modeled, offer stronger predictive power than traditional biomarkers.

Specifically, three key contributions of CRISGI and 128-TER stand out. (1) Automation and scalability. Unlike methods that rely on manual inspection of signal trends, CRISGI enables automated prediction of infection onset and symptom development, providing a scalable pipeline for translational applications. (2) Robust generalization. CRISGI’s predictive power held across different viruses and datasets, emphasizing conserved immune transition mechanisms. This is crucial for real-world deployment where infection sources and cohorts may vary. (3) Biological interpretability. The 128-TER network includes immune-regulatory genes such as IFIT1 and CXCL10, both known to mediate interferon-driven antiviral responses. This not only supports the biological validity of the method but also highlights potential therapeutic targets. However, challenges remain. While CRISGI was developed using bulk transcriptomics, applying it to single-cell data could further refine the cellular resolution. Additionally, prospective validation in large-scale clinical trials will be essential to confirm utility in patient stratification and early intervention.

In conclusion, CRISGI provides a novel lens to explore early, dynamic biomarkers for virus infection and offers a framework adaptable to other diseases governed by complex GRN transitions. The convergence of theory, computation, and validation presented here paves the way for next-generation biomarker discovery.

## Data Availability

The temporal bulk RNA-seq virus infection datasets are downloaded from GEO with accession code GSE30550 (1), GSE52428 (19), GSE73072 (20).

## Author Contribution Statements

CSL: Software; Formal analysis; Validation; Visualization; Investigation; Methodology; Writing—original draft; Writing—review and editing. ANJ: Software; Visualization, Writing—review and editing. KHN: Software. XYL: Investigation. LXC: Conceptualization; Supervision; Software; Visualization; Methodology; Writing—original draft; Writing—review and editing.

